# Structural rearrangements underlying the activation of STIM1 by ER calcium depletion

**DOI:** 10.64898/2026.01.16.700022

**Authors:** Ruoyi Qiu, Richard S. Lewis

## Abstract

In resting cells, STIM1, the dimeric ER Ca^2+^ sensor that controls store-operated Ca^2+^ entry (SOCE), is held in a Ca^2+^-bound inactive state by multiple intramolecular restraints, or brakes. Receptor-evoked release of Ca^2+^ from the ER causes a large conformational change in STIM1 that releases the brakes and exposes the CRAC activation domain (CAD), enabling it to bind and open store-operated Orai1 channels in the plasma membrane. We performed single-molecule FRET measurements with purified STIM1 to better understand how Ca^2+^ release from the luminal domain of STIM1 drives the conformational changes in the cytosolic domain that underlie CAD release. We find that Ca^2+^ removal releases the CAD from CC1α1 (the ‘CC1 clamp’) without obligatory formation of the CC1 coiled-coil that has been associated with CAD release in cells. Surprisingly, the CAD rearranges dramatically during release, as the two hairpin protomers that create its characteristic V-shaped structure are spread apart. Locking the two protomers together by cysteine crosslinking prevents CAD release, suggesting that the CAD must rearrange to escape the CC1 clamp. Our data support a model in which ER depletion-induced dimerization of the luminal SAM domains stabilizes an intermediate 3-helix bundle structure arising from helical interactions of CC1α2 and CC1α3 with CC1α1, thereby releasing the CC1 clamp and allowing the CAD to escape through a ‘fold-out’ mechanism, with subsequent formation of the CC1 coiled-coil enabling the CAD to revert to its original shape to activate Orai1 in vivo.

## INTRODUCTION

Store-operated Ca^2+^ entry (SOCE) is a nearly ubiquitous pathway for Ca^2+^ signaling that underlies many essential physiological processes ranging from gene expression and secretion to immune activation, muscle contraction, and neurological function (Prakriya and Lewis, 2015; Vaeth et al., 2020; Korshunov and Prakriya, 2025). The major site of regulation is the STIM family of ER Ca^2+^ sensors. In mammalian cells, STIM1 senses depletion of Ca^2+^ in the ER and becomes activated to trigger a self-organizing sequence of events in which STIM1 accumulates at ER-plasma membrane (ER-PM) junctions where it binds, traps, and opens Orai1 Ca^2+^ channels in the PM (Prakriya and Lewis, 2015). Because of its central role in regulating SOCE, it is important to understand how STIM1 activity is controlled by ER Ca^2+^ at a structural level. In a recent study, we combined single-molecule FRET (smFRET) measurements with structural modeling using AlphaFold2 to reveal the compact, inactive conformation of STIM1 in the Ca^2+^ bound state, and to identify the intramolecular restraints, or brakes, that maintain it (Qiu and Lewis, 2025). This paper uses a similar approach to describe how Ca^2+^ removal from STIM1 releases these brakes and leads to STIM1 activation.

STIM1 is a dimeric single-pass ER membrane protein, with a luminal domain that senses Ca^2+^ and multiple cytosolic domains that control interactions with the PM and Orai1 (**Fig. 1A**). The critical part of STIM1 that controls Orai1 activity is the CRAC activation domain (CAD; also known as SOAR or Ccb9) (Park et al., 2009; Yuan et al., 2009; Kawasaki et al., 2009). In resting cells, the luminal ER [Ca^2+^] of ∼0.5 mM saturates the EF-hands, which bind to the luminal SAM domains and maintain the cytosolic domain in an inactive conformation (Stathopulos et al., 2006, 2008; Gudlur et al., 2018). Under these resting conditions, multiple weak brakes hold the apex of CAD against the ER membrane (**Fig. 1C**); these include a steric restraint from the Ca^2+^-bound EF-SAM, a domain-swapped hydrophobic interaction of CC1α1 with CC3 of CAD (often referred to as the ‘CC1 clamp’), intersubunit hydrophobic and electrostatic interactions between the CC1α2 and CC1α3 domains, and electrostatic interactions of the CAD apex with charged lipids in the ER membrane (Fahrner et al., 2014; Ma et al., 2015; Qiu and Lewis, 2025).

**Figure 1.**
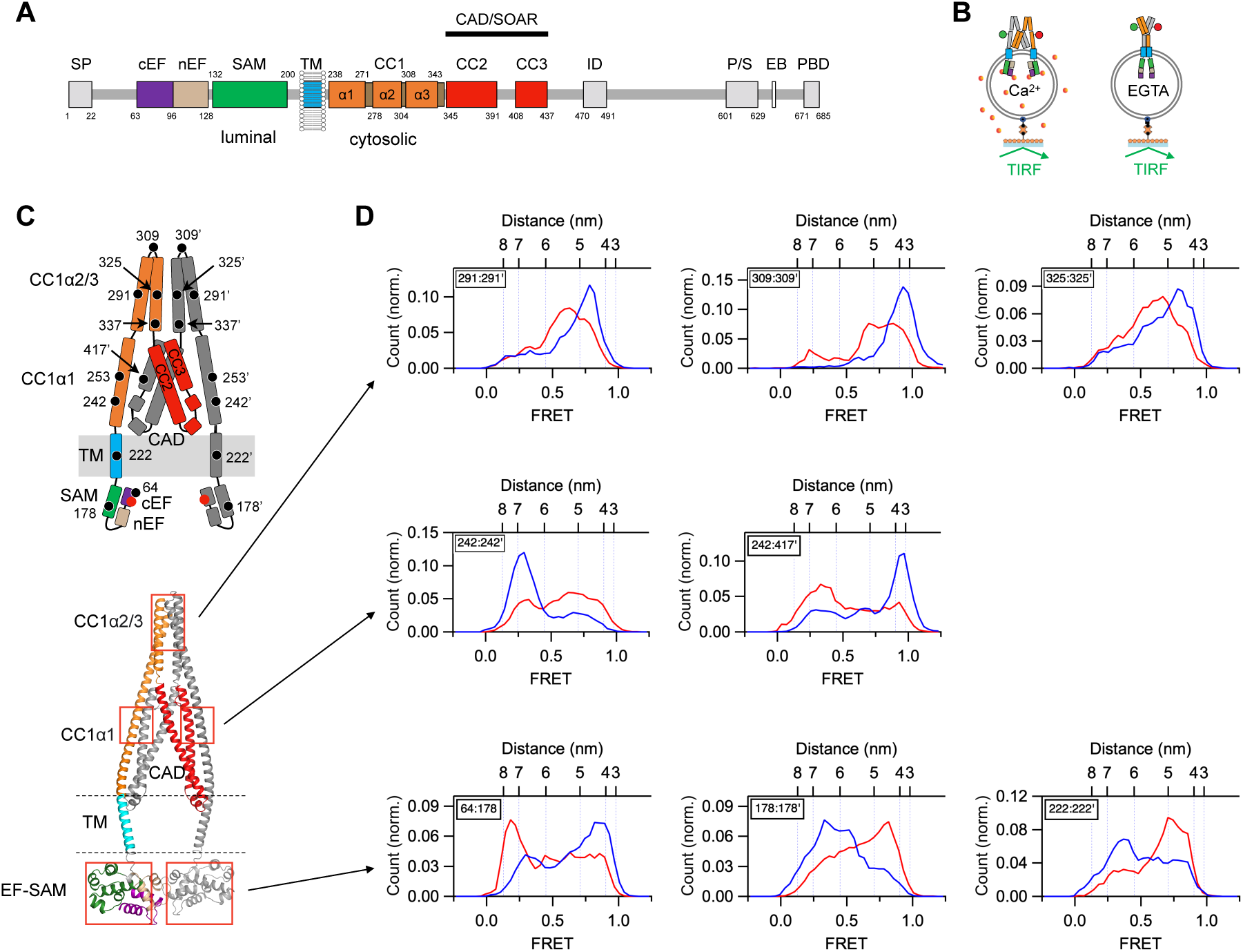
EGTA releases multiple brakes throughout STIM1. (*A*) Domain organization of flSTIM1. SP, signal pepMde; cEF, canonical EF hand; nEF, non-canonical EF hand; SAM, sterile alpha moMf; TM, transmembrane domain; CC1-3, coiled-coil 1-3; ID, inacMvaMon domain; P/S, proline/serine-rich region; EB, EB1 binding domain; PBD, polybasic domain; CAD, CRAC acMvaMon domain; SOAR, STIM-Orai acMvaMon region. Residue numbers for each domain are indicated. (*B*) flSTIM1 reconstituted into liposomes in presence of 2 mM Ca^2+^ or 0.5 mM EGTA and attached to coverslips. (*C*) Schematic diagram (*Top*) and AlphaFold2 model (*Bottom*) of the resting state of flSTIM1, with boxes showing the location of the EF-SAM, CC1α1-CC3, and CC1α2/3 brakes. (*D*) smFRET histograms in 2 mM Ca^2+^ (blue) or EGTA (red) are shown for the indicated dye pairs. EGTA releases the EF hand from SAM (64:178) and brings the SAM domains (178:178’) and TM domains (222:222’) closer together (*Bottom*). EGTA destabilizes the CC1α1-CC3 clamp and releases CAD (242:242’ and 242:417’) (*Middle)*. EGTA releases the CC1α2/3 brake, shown by increasing the intersubunit distances between the CC1α2 domains (291:291’), the CC1α2-3 linkers (309:309’) and the CC1α3 domains (325:325’) (*Top*). Numbers of molecules (in Ca^2+^, in EGTA): 64:178 (n=213, 233); 178:178’ (n=271, 282); 222:222’ (n=114, 83); 242:242’ (n=361, 437); 242:417’ (n=190, 224); 291:291’ (n=185, 182); 309:309’ (n=76, 88); 325:325’ (n=243, 251).

Upon agonist-induced release of Ca^2+^ from the ER, the decline of [Ca^2+^]_ER_ below ∼200 µM evokes a series of STIM1 conformational changes. Ca^2+^ unbinding from the EF hand exposes and rearranges the SAM domains, promoting their dimerization (Stathopulos et al., 2006; Gudlur et al., 2018), and leading to coiled-coil formation by the TM and proximal CC1α1 domains (Hirve et al., 2018). These rearrangements are in turn linked to the release of the C-terminal polybasic domain (PBD) that enables STIM1 trapping at ER-PM junctions, and to the release of the CAD from the CC1 clamp, followed by a 180° reorientation and translocation of CAD to bind and activate Orai1 across the 10-15 nm cleft of the ER-PM junction (Prakriya and Lewis, 2015). Remarkably, all of these conformational rearrangements are reversible, as STIM1 reverts to its resting state upon refilling of the ER lumen.

The reversible remodeling of STIM1 during activation occurs globally throughout the cytosolic domain. Many essential aspects of the process remain undefined, in large part due to the difficulty of applying approaches like x-ray crystallography or cryo-EM to obtain structures of this highly flexible protein. In this study, we used smFRET (Roy et al., 2008) to describe the conformations and dynamics of STIM1 under Ca^2+^-free conditions in order to understand the mechanisms underlying STIM1 activation. The results reveal structural rearrangements of the EF-SAM luminal domain and the CC1 and CAD cytosolic domains that are required for the release of CAD from the CC1 clamp, as well as potential intermediates in this process. We propose a model in which structural fluctuations throughout STIM1 enable SAM dimerization to trap the cytosolic domains in conformations that lead to CAD restructuring and release. While coiled-coil formation is not required for CAD release, it may serve to refold the CAD prior to binding and activation of Orai1.

## RESULTS

### Ca^2+^ removal releases multiple brakes to drive STIM1 towards the activated state

After removing all endogenous cysteines in full-length STIM1 (flSTIM1), we introduced pairs of cysteines, labeled them with donor and acceptor dyes to serve as smFRET probes, and reconstituted the labeled protein in liposomes prior to attachment to coverslips for TIRF microscopy (**Fig. 1B**). Using this approach, we have previously shown that multiple intramolecular interactions create four restraints, or brakes, that stabilize the inactive state of Ca^2+^-bound STIM1: the EF-SAM domain, the CC1α1-CC3 clamp, CAD apex-lipid attraction, and the CC1α2/3 dimer (**Fig. 1C**) (Qiu and Lewis, 2025). Removal of Ca^2+^ with EGTA drives major rearrangements throughout the entire STIM1 structure that release all four brakes. (Throughout this paper, we use ‘:’ to denote sites in the dimer, and ‘’’ to indicate a site on the partner subunit.) In smFRET histograms, shifts in the predominant FRET level show that as the EF hand is released from the SAM domain (64:185), the two SAM domains (178:178’) and two TM domains (222:222’) come closer together (**Fig. 1D**, bottom row). The interaction between CC1α1 and CC3’ in CAD is broken, releasing CAD from the CC1α1 brake (242:417’) and allowing the CC1α1 helices to come closer together (242:242’) (**Fig. 1D**, middle row). Finally, the CC1α2/3 domains separate (291:291’, 309:309’, 325:325’), releasing the CC1α2/3 brake (**Fig. 1D**, top row). These results underscore the global rearrangements throughout STIM1 driven by Ca^2+^ release from the luminal domain.

### Ca^2+^ removal releases CAD without formation of a CC1 coiled-coil

Based on previous studies of STIM1 in cells, we expected to find that under Ca^2+^-free conditions STIM1 would adopt a state in which the TM and CC1α1 domains form a dimeric coiled-coil (Hirve et al., 2018). The smFRET histogram for 242:242’ in EGTA can be fitted by a sum of three Gaussian components the highest of which, centered at a FRET level of 0.87, corresponds to a distance of 4.2 nm (**Fig. 2A**). Measurement between the TM domains at 222:222’ gave a similar maximum FRET value of 0.86 (4.3 nm; **Fig. 2B**). These distances are significantly greater than expected for dye-dye distances on either side of a symmetric coiled-coil, which we estimated to be ∼3.3 nm using cortexillin I as a canonical 2-stranded coiled-coil (**Fig. S1**). To drive STIM1 towards the coiled-coil state we introduced the double mutation N234L/S237L (STIM1-LL), which promotes spontaneous coiled-coil formation in the TM and CC1α1 domains and constitutively activates SOCE in cells by overriding a discontinuity in the heptad repeat between the two domains (Hirve et al., 2018). The peak FRET for this mutant in EGTA corresponds to distances of 3.1 nm for 242:242’ and 3.3 nm for 222:222’, close to that predicted for cortexillin I (**Fig. 2C, D**). In addition, the CC1α2/3 hairpin structure unfolds and extends, as indicated by a large increase in the distance between 291 and 325 within each CAD protomer (**Fig. S2**). These results confirm the ability of the N234L/S237L mutation to promote TM-CC1α1 coiled-coil formation and demonstrate that in the WT protein this structure fails to form in EGTA.

**Figure 2.**
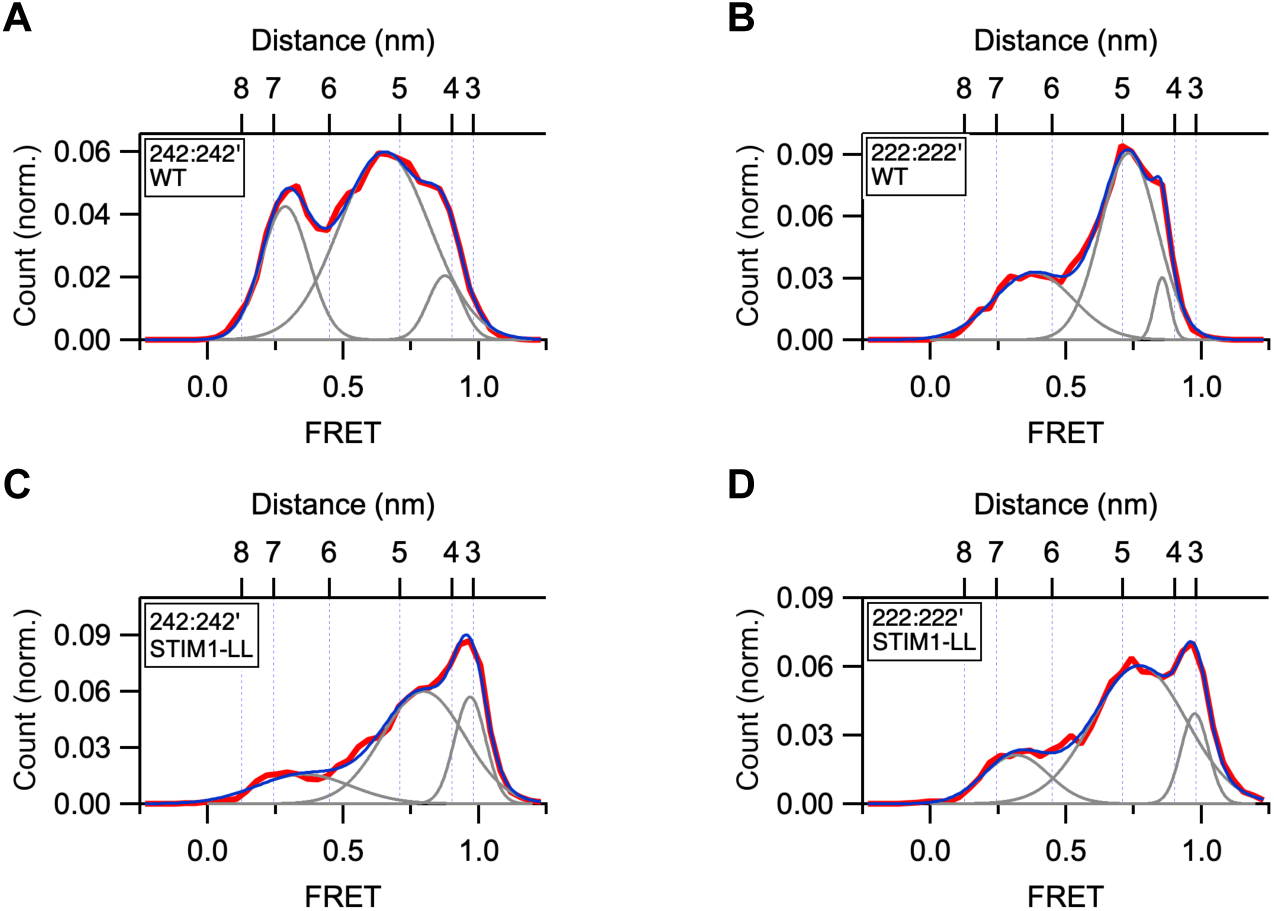
EGTA fails to drive the CC1 domains in WT STIM1 to form a coiled-coil. All FRET distributions were obtained in 0.5 mM EGTA. FiÄed Gaussian curves (gray) and their sum (blue) are superimposed on the data (red). (*A*) CC1α1-CC1α1’ FRET histogram of WT STIM1 (242:242’; n=437). Fit parameters (peak FRET and fracMonal area): 0.29 (25%), 0.65 (67%), 0.88 (8%). The smallest distance (4.2 nm) is greater than expected for a coiled-coil. (*B*) TM-TM’ FRET histogram of WT STIM1 (222:222’; n=83). Fit parameters (peak FRET and fracMonal area): 0.38 (32%), 0.73 (62%), 0.86 (6%). The minimum TM-TM’ distance (4.3 nm) is greater than expected for a coiled-coil. (*C*) CC1α1-CC1α1’ FRET histogram of STIM1-LL (242:242’; n=254). Fit parameters (peak FRET and fracMonal area): 0.36 (18%), 0.80 (60%), 0.97 (22%). The smallest distance (3.3 nm) is consistent with a coiled-coil. (*D*) TM-TM’ FRET histogram of STIM1-LL (222:222’; n=203). Fit parameters (peak FRET and fracMonal area): 0.32 (17%), 0.77 (70%), 0.98 (13%). The minimum TM-TM’ distance (3.1 nm) is consistent with a coiled-coil.

In light of these results, we considered the possibility that the CC1α1, 2, and 3 domains might rearrange in EGTA to form a 3-helix bundle (3HB) structure that was described by NMR spectroscopy for the CC1 monomer in solution (pdb: 6YEL) (Rathner et al., 2020). In this structure, CC1α2 binds to the same residues in CC1α1 that form the CC1-CC3 clamp, suggesting that competition for CC1α1 may enable the 3HB to disrupt the clamp. We selected four intramolecular dye pair locations within CC1 to test for the 3HB structure: 242:291, 242:309, 253:291, and 291:325. With the exception of 291:325, these dye pairs are well separated in 2 mM Ca^2+^, consistent with the resting STIM1 structure, and come closer together in EGTA (**Fig. S3**). As shown in **Fig. 3**, in EGTA the distances estimated from the highest FRET components of the histograms agree well with the simulated dye distances from the NMR structure, consistent with a fraction of STIM1 molecules in the 3HB configuration in the steady-state. In addition, an intermediate FRET component with a 51-67% probability is present for each dye pair, possibly representing a looser arrangement of the 3HB.

**Figure 3.**
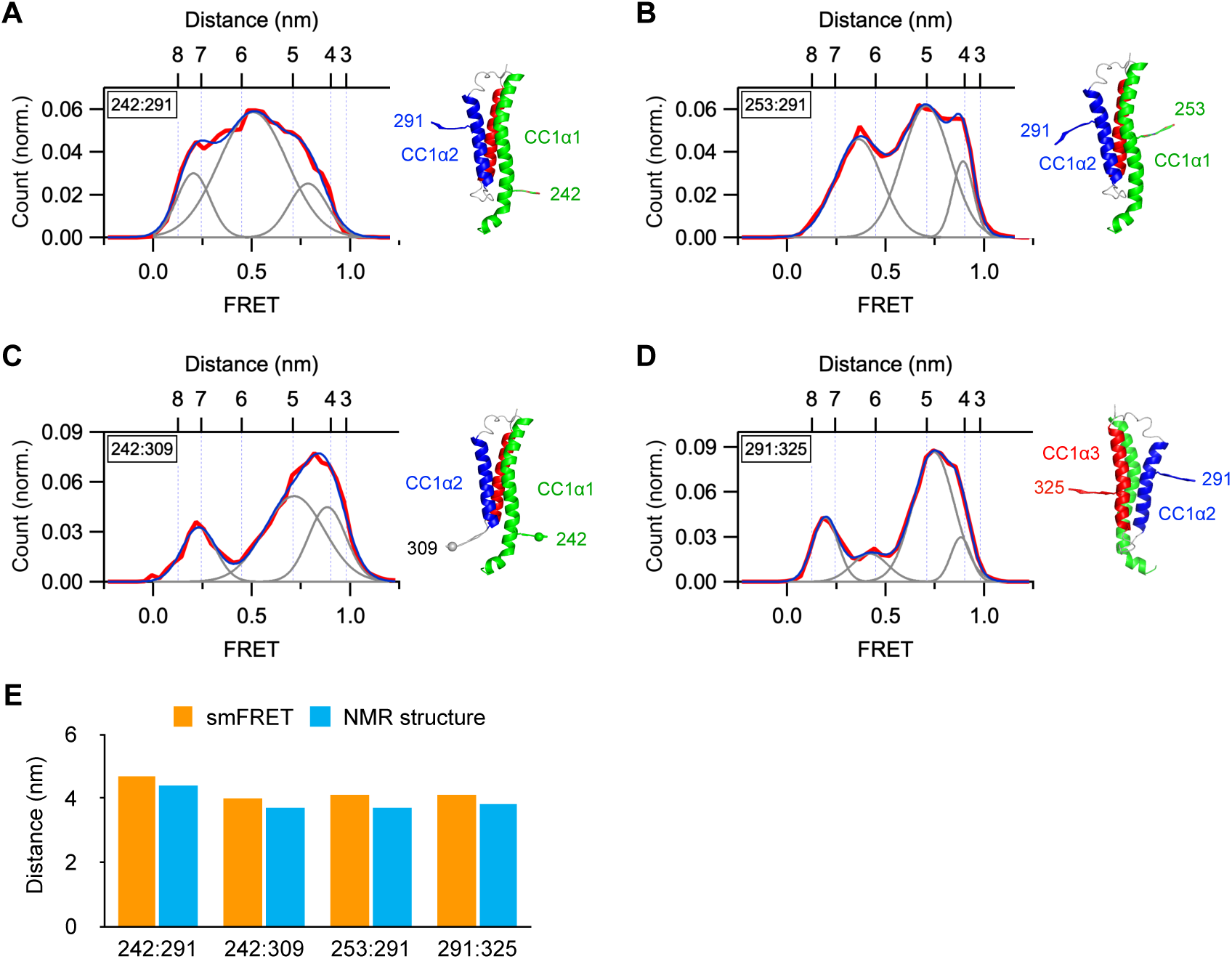
The CC1 domains form a 3-helix bundle structure in EGTA. All FRET measurements were obtained for WT STIM1 in 0.5 mM EGTA. In each panel, simulated dye positions are superimposed on the 3HB structures (pdb: 6YEL). (*A*) smFRET histogram for 242:291 (n=189). Fit parameters (peak FRET and fracMonal area): 0.21 (17%), 0.51 (67%), 0.79 (16%). (*B*) smFRET histogram for 253:291 (n=275). Fit parameters (peak FRET and fracMonal area): 0.36 (36%), 0.71 (51%), 0.89 (13%). (*C*) smFRET histogram for 242:309 (n=158). Fit parameters (peak FRET and fracMonal area): 0.23 (19%), 0.72 (52%), 0.89 (29%). (*D*) smFRET histogram for 291:325 (n=210). Fit parameters (peak FRET and fracMonal area): 0.19 (18%), 0.42 (11%), 0.75 (60%), 0.88 (10%). (*E*) Comparison of the minimal inter-dye distances measured from smFRET and the NMR-derived 3HB structure of monomeric CC1. Intrasubunit distances within CC1 are consistent with formation of the 3-helix bundle.

As further tests of the 3HB structure, we introduced mutations that were expected to destabilize it. R304W, a naturally occurring mutation in the CC1α2-CC1α3 linker, generates constitutive STIM1 activity and SOCE, leading to Stormorken’s syndrome in human patients (Nesin et al., 2014; Lacruz and Feske, 2015). The mutation is thought to activate STIM1 by promoting helix extension through the CC1α2/3 linker and destabilizing the 3HB (Fahrner et al., 2018; Rathner et al., 2020). In the presence of Ca^2+^, this mutation greatly reduced the resting state FRET measured for 242:242’ and generated a high FRET peak corresponding to a distance of 3.6 nm and whose probability increased in EGTA, suggesting that a coiled-coil structure may have formed (**Fig. S4A, B**). It is possible that the high activity of the R304W mutant derives from its ability to destabilize the resting CC1α2/3 brake as well as the 3-helix bundle state, promoting a transition from the resting to the fully active coiled-coil structure. We also examined mutations of I290 and A293, which interact with L258 and L261 to help stabilize the 3HB (Rathner et al., 2020). Like the R304W mutant, the I290S/A293S double mutation also increased the proportion of molecules reaching a high FRET state consistent with coiled-coil formation in EGTA (**Fig. S4C, D**). Together, the effects of these CC1 mutations support the notion of the 3-helix bundle as an intermediate between the inactive and coiled-coil states, such that destabilizing this structure favors a more direct transition to the coiled-coil state (see Discussion).

### Locking the CC1α2/3 brake prevents release of the CC1 clamp and CAD in EGTA

For STIM1 to be able to bind and activate Orai1 after store depletion, the CAD must first be released from the CC1 clamp. We have shown that the CC1α2/3 dimer in Ca^2+^-bound STIM1 acts as a brake to inhibit spontaneous release of CAD in resting cells (Qiu and Lewis, 2025). EGTA releases the CC1α2/3 brake, as shown by a several-nm increase in the separation of the S337 residues at the C-termini of the CC1α3 helices (**Fig. 4A**). In a previous study we found that cysteines substituted for S339 in flSTIM1 in cells and in ctSTIM1 (the cytosolic STIM1 233-685 fragment) in vitro could be disulfide-crosslinked with the oxidizing agent diamide (van Dorp et al., 2021). Exposure of cells to 1 mM diamide for 10 min crosslinked 64 % of STIM1-S339C, and this was reversed by in vitro treatment with the reducing agent DTT (**Fig. 4B**; see Methods). Based on our AlphaFold2 model of the STIM1 resting state (**Fig. 1C**) (Qiu and Lewis, 2025), crosslinking S339C would be expected to prevent the two hairpins of the CC1α2/3 brake from separating, allowing us to test whether release of the brake is required for CAD release from the CC1α1 clamp. Crosslinking S339C in MSP2N2 nanodiscs with diamide effectively prevented the release of CAD from the CC1α1 clamp in EGTA as monitored by the 242:242’ distance (**Fig. 4C, D**). This inhibition was due to disulfide bond formation, as normal release was restored by DTT (**Fig. 4E**). These results confirm that release of the CC1α2/3 brake is necessary before the CAD can be released from the CC1α1 clamp. We also tested the effect of crosslinking STIM1-S339C on the accumulation of STIM1 at ER-PM junctions in cells following store depletion with thapsigargin (TG). In HEK 293 cells lacking Orai (Orai1/2/3 triple-knockout) (Yoast et al., 2020), diamide pretreatment slowed the formation of STIM1-S339C puncta, though the final extent of accumulation was unaffected (**Fig. 4F, Fig. S5**). These results may be explained by the reversibility of disulfide bond formation in the reducing environment of the cytoplasm; after store depletion, this would be expected to produce an exponential decay in the disulfide form and a concomitant increase in CAD and PBD release over time, thereby producing an apparent delay in the diffusional trapping of STIM1 at ER-PM junctions. Because TG keeps the stores depleted, STIM1 activation is irreversible, and the mutant STIM1 accumulates in puncta to the same steady-state level as WT.

**Figure 4.**
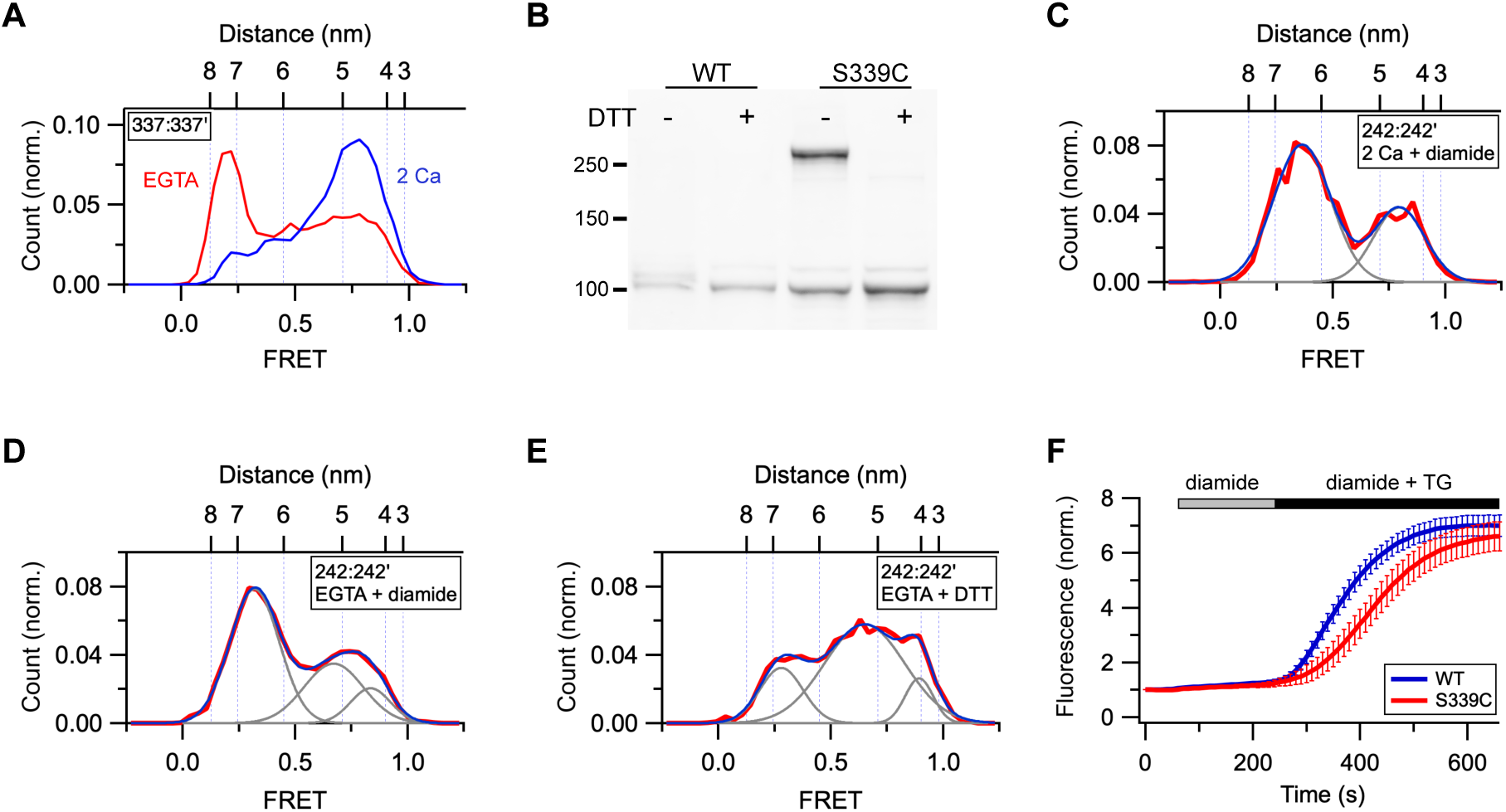
Locking the CC1α2/3 brake prevents the release of CAD in EGTA. (*A*) EGTA greatly increases the separation between CC1α3-CC1α3’ (337:337’). (*B*) Western blot shows diamide treatment of STIM1-S339C expressed in HEK 293 cells induces disulfide-linked dimers that are sensitive to DTT. Molecular weights in kDa are indicated. (*C-E*) S339C crosslinking prevents release of CAD from the CC1α1 clamp. (*C*) smFRET histogram for 242:242’ in 2 mM Ca^2+^ + diamide (n=69). Fit parameters (peak FRET and fracMonal area): 0.36 (67%), 0.80 (33%). (*D*) smFRET histogram for 242:242’ in EGTA + diamide (n=229). Fit parameters (peak FRET and fracMonal area): 0.32 (58%), 0.67 (31%), 0.84 (12%). (*E*) smFRET histogram for 242:242’ in EGTA following DTT treatment (n=210). Fit parameters (peak FRET and fracMonal area): 0.28 (21%), 0.65 (69%), 0.89 (11%). (*F*) Effect of diamide treatment on puncta formation by STIM1-S339C following store depletion with TG. Mean (± sem) of 27 (WT) or 15 (S339C) cells.

### Energetic coupling of cytosolic and luminal domains of STIM1 is bidirectional

The multiple conformational changes throughout the cytosolic domain evoked by Ca^2+^ release from the cEF hand is an example of anterograde coupling between luminal and cytosolic domains across the ER membrane (**Fig. 1**). Retrograde coupling is also evident from the effects of activating and inactivating mutations in the cytosolic domain on the conformation of the luminal domain. For the 178:178’ dye pair in the SAM domain, EGTA shifts FRET to higher values, indicating closer apposition of the two SAM domains in the Ca^2+^-free state (**Fig. 1D**). In the presence of saturating Ca^2+^, introduction of the activating N234L/S237L mutation also shifts 178:178’ FRET to higher values (**Fig. 5A**). EGTA further augmented the FRET, presumably by increasing the exposure of the SAM dimerization interface (**Fig. 5B**). Conversely, crosslinking the S339C mutant with diamide to inhibit CAD release in EGTA diminished the high FRET (**Fig. 5C**). These results show that energetic coupling between luminal and cytosolic domains of STIM1 is bidirectional, consistent with a thermally-driven passive rather than active (e.g., ATP-dependent) mechanism of activation.

**Figure 5.**
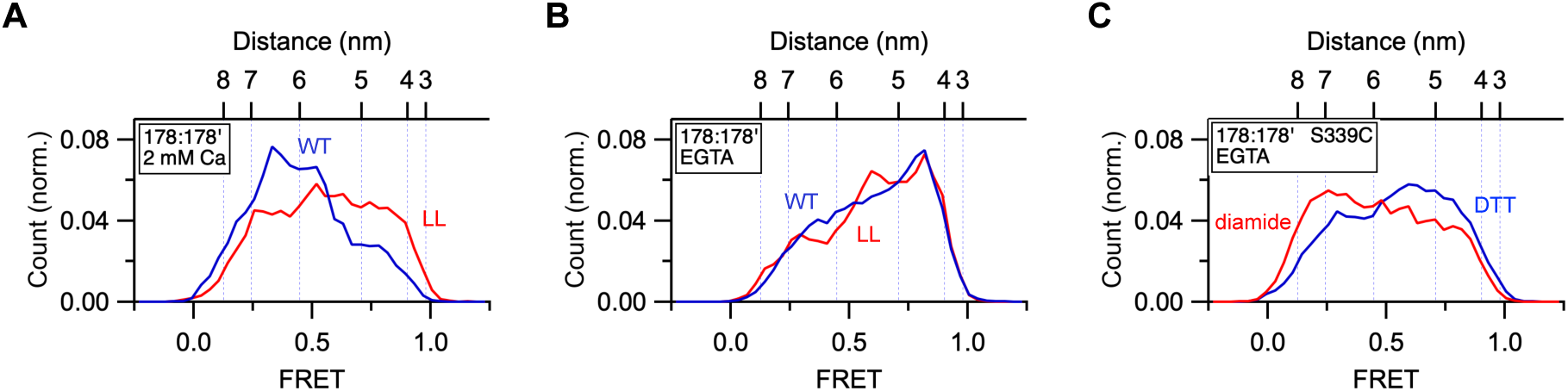
Retrograde energetic coupling of cytosolic and luminal domains of STIM1. (*A*) 178:178’ FRET in the presence of saturating Ca^2+^ for WT STIM1 (blue; n=271)) and STIM1-LL (red; n=253). Promotion of the CC1 coiled-coil moves the SAM domains closer together despite the presence of Ca^2+^. (*B*) 178:178’ FRET in the presence of EGTA for WT STIM1 (blue; n=282) and STIM1-LL (red; n=189). (*C*) 178:178’ FRET in EGTA for STIM1-339C in the presence of diamide (blue; n=371) or DTT (red: n=206). Locking the CC1α2/3 brake impedes the movement of SAM domains in EGTA.

### Ca^2+^ removal changes the structure of the CAD

One way in which CAD could be released from the CC1 clamp is through a simple ‘flip-out’ mechanism in which it retains its V-shaped structure and moves as a rigid body (van Dorp et al., 2021). To test this possibility, we measured distances at multiple sites within CAD in the presence of Ca^2+^ or EGTA. In Ca^2+^, predominant high FRET peaks were evident at sites between the N-termini (349:349’), apices (378:378’), and C-termini (431:431’) of the two CAD protomers, as well as between the N– and C-termini of each protomer (349:431); the corresponding smFRET-derived distances were consistent with the close packing of the two hairpin-shaped subunits in the CAD crystal structure (Yang et al., 2012) (**Fig. 6A-D** blue histograms and **Table S1**). EGTA significantly dilated the CAD structure. The FRET for 378:378’ was shifted towards lower values, indicating a separation of the two apical regions of CAD in EGTA (**Fig. 6B**). In addition, the distances between the CAD N-termini (349:349’) and between the C-termini (431:431’) both increased, showing that the two CAD monomers move apart in response to Ca^2+^ removal (**Fig. 6A** and **C**). To test whether these changes reflect a simple movement of the two CC2-CC3 hairpin structures as rigid bodies, we measured the distance between the N– and C-termini within a protomer using the 349:431 dye pair (**Fig. 6D**). The reduction of the high-FRET population and emergence of a low-FRET peak shows that the CC2 and CC3 domains of each hairpin also separate from one another in EGTA. Interestingly, the distance between the N-terminus of one CAD protomer with the C-terminus of the other (349:431’) did not change (**Fig. 6E**), suggesting that the two monomers remain connected at their base. Taken together, the results suggest that CAD structure dilates as the C/N’ and C’/N ends are pulled apart (**Fig. 6F**). These results reveal that the CAD structure in STIM1 is not rigid but is modified under the Ca^2+^-free conditions that release the CC1 clamp. The R304W mutation, which promotes CC1 coiled-coil formation in EGTA (**Fig. S4B**), restored the high-FRET peak for 431:431’, suggesting that the released CAD reverts to its resting structure after CC1 rearranges into a coiled-coil (**Fig. 6C**).

**Figure 6.**
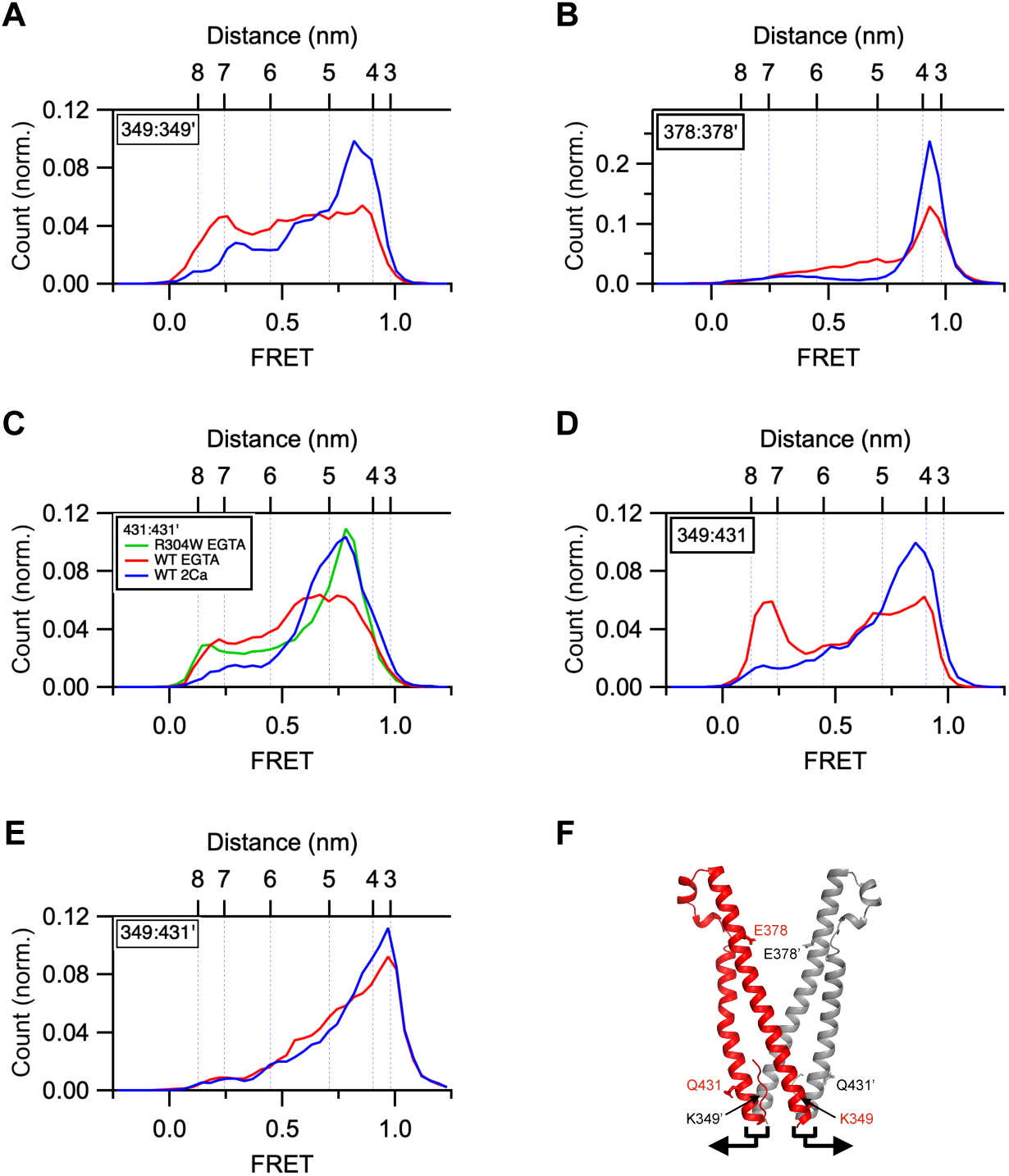
EGTA treatment rearranges the structure of the CAD in flSTIM1. FRET measurements were obtained in 2 mM Ca^2+^ (blue) or 0.5 mM EGTA (red). (*A*) Distance between the N-termini (349:349’) of the two CAD protomers increases in EGTA; n=221 (Ca^2+^), n=247 (EGTA). (*B*) Distance between the CAD apices (378:378’) increases in EGTA; n=226 (Ca^2+^), n=219 (EGTA). (*C*) Distance between the C-termini (431:431’) increases in EGTA; n=204 (Ca^2+^), n=213 (EGTA). R304W restores the high FRET peak in Ca^2+^; n=290. (*D*) The N– and C-termini of each CAD protomer (349:431) separate in EGTA; n=223 (Ca^2+^), n=221 (EGTA). (*E*) The N-terminus of each protomer stays associated with the C terminus of its neighbor (349:431’) in EGTA; n=232 (Ca^2+^), n=186 (EGTA). (*F*) Crystal structure of human STIM1 CAD showing the dye locations in A-E (3TEQ.pdb). Arrows illustrate movement of the paired N and C termini of the two protomers in EGTA consistent with distance meaurements in A-E.

### Locking the two CAD protomers together prevents CAD rearrangement and the release of the CC1 clamp

While the results clearly indicate structural rearrangments of the CAD in EGTA, they do not show whether these are required for release of the CC1 clamp. To address this question, we sought a way to lock the two CAD protomers together in STIM1. Based on the CAD crystal structure, the two Y361 side chains at the CC2 crossing point are known to engage in a π-π interaction that helps stabilize the CAD dimer (Yang et al., 2012). We found upon substituting cysteine for Y361 that STIM1 could be efficiently crosslinked with diamide (84%), and this was effectively reversed by treatment with DTT (**Fig. 7A**). Based on these results, we treated purified STIM1-Y361C in MSP2N2 nanodiscs with diamide and conducted smFRET measurements in the presence of 2 mM Ca^2+^ or 0.5 mM EGTA.

**Figure 7.**
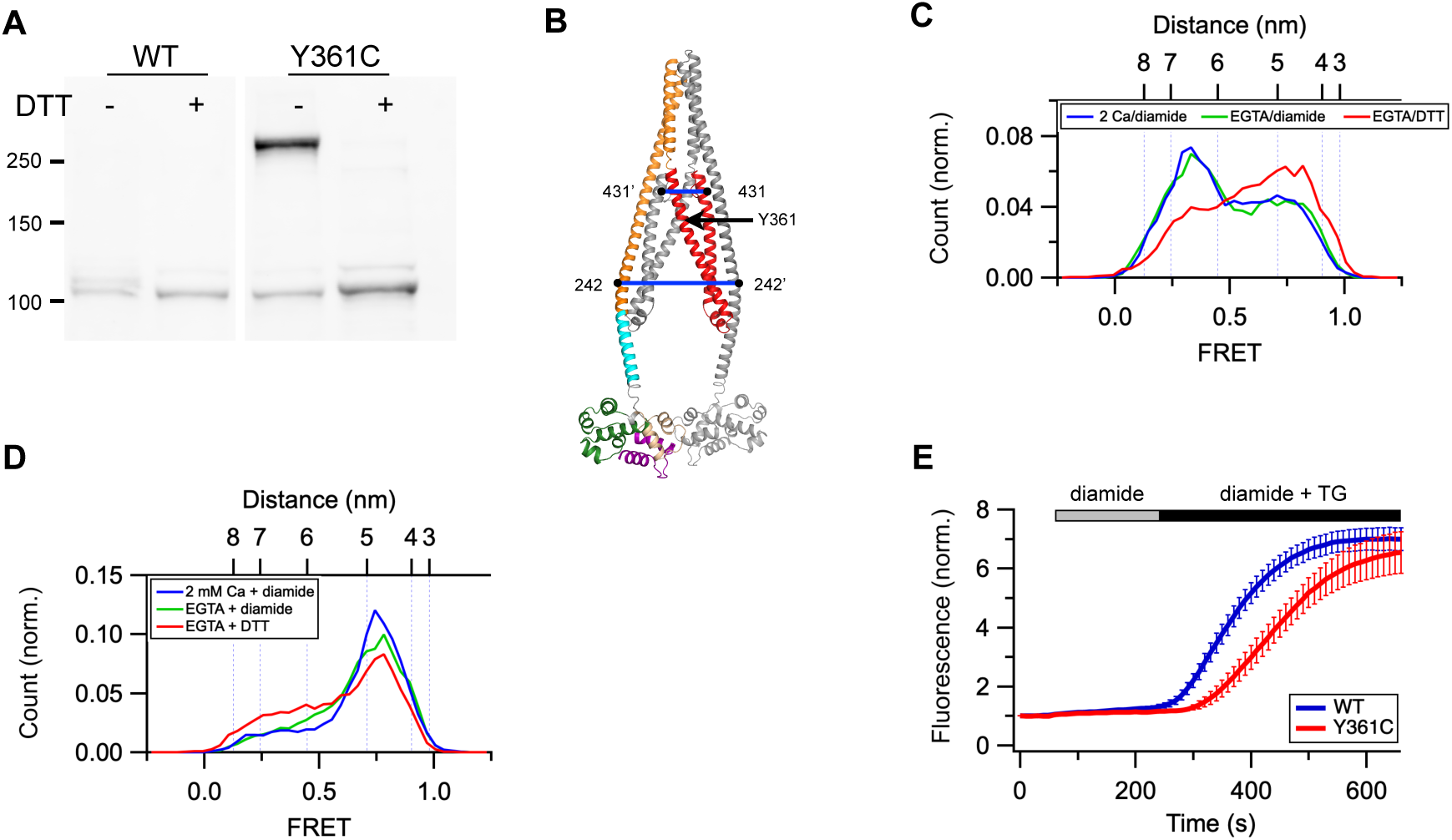
Locking the CAD structure prevents its escape from the CC1α1 clamp in EGTA. (*A*) Western blot shows diamide treatment of STIM1-S361C expressed in HEK 293 cells induces disulfide-linked dimers that are sensitive to DTT. Molecular weights in kDa are indicated. (*B*) AlphaFold2 model of the STIM1 resting state showing the Y361C crosslinking site and dye locations for FRET measurements. (*C*) smFRET histograms for 242:242’ in STIM1-Y361C in 2 mM Ca^2+^ + diamide (blue), EGTA + diamide (green), and EGTA + DTT (red). (*D*) smFRET histograms for 431:431’ in STIM1-Y361C in 2 mM Ca^2+^ +diamide (blue), EGTA + diamide (green), and EGTA + DTT (red). (*E*) Effect of diamide treatment on puncta formation by STIM1-Y361C following store depletion with TG. Mean (± sem) of 27 (WT) or 17 (Y361C) cells.

The Y361C mutation by itself appeared to destabilize the Ca^2+^-bound STIM1 resting state slightly, as judged by the reduced occupancy of the low-FRET state for 242:242’ (cf. **Fig. 7C** and **Fig. 1E**). This effect may be a consequence of eliminating the Y361 π-π interactions that stabilize the CAD dimer. Importantly, diamide treatment prevented CAD release in EGTA, and release was rescued by the addition of DTT to reduce the disulfide bonds (**Fig. 7C**, green and red histograms). Similarly, diamide treatment to crosslink Y361C prevented the separation of CAD C-termini (431:431’) in EGTA, and this was also reversed by DTT (**Fig. 7D**, green and red histograms). Thus, locking the CAD protomers together at the Y361 crossover point inhibits the separation of the CAD C-termini and release of the CC1 clamp just as effectively as does locking the CC1α2/3 brake (**Figs. 4** and **7**). As we observed for the S339C mutant, diamide-induced crosslinking of STIM1-Y361C significantly slowed the formation of puncta in response to store depletion, as expected from the effect of crosslinking on release of the CC1 clamp (**Figs. 7E** and **S5**).

## DISCUSSION

In response to store depletion, STIM1 undergoes a massive conformational change, rearranging from an inactive state in which CAD is held against the ER membrane, to an active state where it is rotated and extended across the 15-nm gap at ER-PM junctions to bind and activate Orai1. From a series of smFRET-based distance measurements of purified STIM1 in vitro, we have described critical rearrangements of the cytosolic domains that reveal potential intermediates in the activation process and suggest a structural pathway that couples Ca^2+^ release from the luminal domain to STIM1 activation, while raising new questions about the active state structures of STIM1 in cells.

Previous work by Hogan and colleagues (Hirve et al., 2018) has shown that Ca^2+^ removal from STIM1 in native ER membranes causes the TM and CC1α1 domains to form an extended coiled-coil, implicating the coiled-coil in the release of CAD from the CC1 clamp to activate Orai1 and SOCE. In our studies of reconstituted WT STIM1 in vitro, we were surprised to find that CAD was released in EGTA despite an apparent lack of coiled-coil formation in the TM and CC1α1 domains based on intersubunit distances (**Fig. 2**). The lack of coiled-coils could be related to the absence of multiple proteins and conditions in our purified protein system that are thought to enhance the efficiency of STIM1 activation in vivo. One such candidate is the ER membrane protein STIMATE (TMEM110), a strong positive modulator of SOCE. STIMATE has been shown to bind the CC1 domain and inhibit CC1-CAD binding (Jing et al., 2015), suggesting that it stabilizes the extended state of STIM1. Other STIM1-interacting proteins such as CRACR2A (Srikanth et al., 2010), as well as interaction of STIM1 with PIP_2_ and PI4P in the plasma membrane (Bhardwaj et al., 2013; Cohen et al., 2023), oligomerization of STIM1 (Luik et al., 2008; Liou et al., 2007), and physiological temperature (37°C) (Yarotskyy and Dirksen, 2012; Xiao et al., 2011) have all been associated to varying degrees with an increase in activation and could also increase the stability of the coiled-coil state directly or indirectly by stabilizing the SAM-SAM dimer.

Instead of a coiled-coil, we found that the CC1 domain rearranged into multiple conformations, including a 3-helix bundle structure similar to that of purified monomeric CC1 protein in solution (Rathner et al., 2020) (**Fig. 3**). Rathner et al showed that the same CC1α1 residues that make the CC1-CC3 clamp (L248, L251, L258, and L261) are engaged in coiled-coil interaction with CC1α2 in the 3HB, and mutations predicted to weaken this interaction in the 3HB, such as I290S/A293S, slowed I_CRAC_ induction without affecting its amplitude. To explain these results, the authors proposed that the 3HB competes with the CC1α1-CC3 pair, such that weakening the 3HB strengthens the CC1α1-CC3 clamp. In a similar way, we believe the 3HB may compete with the CC1 coiled-coil, explaining why the R304W and I290S/A293S mutations, both of which destabilize the 3HB, increase the steady-state fraction of molecules that form the coiled-coil state in EGTA (**Fig. S4**). As an intermediate between the resting and coiled-coil states, the 3HB may serve two functions. First, through hydrophobic interactions the 3HB could mitigate the energetic cost of exposing hydrophobic CC1α1 residues to the aqueous environment when the CC1-CC3 brake is released and the coiled-coil has not yet formed. Similarly, the 3HB provides a favorable environment for the hydrophobic residues (V324, L335) that would be exposed upon release of the CC1α2/3 brake (Qiu and Lewis, 2025). In addition, by providing a low-energy intermediate, the 3HB may moderate the formation of the coiled-coil to ensure the ready reversibility of CC1 conformational changes upon refilling of the Ca^2+^ stores. A similar moderating role was first recognized for the sentinel residues N234 and S237 that weaken the CC1α1 coiled-coil (Hirve et al., 2018).

During STIM1 activation, CAD is freed from its interaction with CC1α1, rotates, and is extended towards Orai1 in the PM. We have proposed two possible ways in which CAD could be released from the CC1 clamp: a simple ‘flip-out’ model in which the CAD V-shaped structure is maintained and moves as a rigid body, or a more complex ‘fold-out’ model in which some refolding of the CAD dimer occurs to allow release (van Dorp et al., 2021). We observed significant restructuring of the CAD in EGTA, consistent with a movement of the paired N/C’ and N’/C termini of the two CAD protomers away from each other during release (**Fig. 6F**). Maintained association of these terminal domains is consistent with a network of hydrophobic interactions and H bonds in the CAD crystal structure between the N terminus of each protomer with the C terminus of its partner (Yang et al., 2012). This restructuring is not simply a result of release from the CC1 clamp, as the CAD base is not detectable affected by removing CC1 from the soluble ctSTIM1 fragment (van Dorp et al., 2021). Instead, it seems dependent on rearrangements of CC1 that occur in flSTIM1 in EGTA. Most importantly, the unfolding of CAD is required for release from the CC1α1 clamp, as locking the CAD structure by crosslinking the two protomers at Y361C effectively prevents release in the presence of EGTA (**Fig. 7**). These results argue against a flip-out model in which CAD retains its structure throughout the release process, but rather support some kind of CAD restructuring as represented in the fold-out model. This conclusion is consistent with molecular dynamics simulations of a STIM1 model in which the two CC2 domains move spontaneously in opposite directions (Horvath et al., 2023). Our results suggest that the CAD base reverts to its close-packed structure when the two CC1α1 domains fold into a coiled-coil (**Fig. 6C**). One possibility is that the two N-termini of CAD, which are attached to the CC1α3 helices, are held apart by the bulk of the 3HBs, and that disassembly of the 3HB allows the CAD to revert to its lowest energy conformation with a closely packed basal domain. Several important questions related to the coiled-coil state arise from these results. To what extent might the 3HB state be active in cells; can it release the PBD for accumulation at ER-PM junctions, and could CAD reach Orai1 without formation of the coiled-coil? Interestingly, in one study deletion of CC1α2 and CC1α3 from STIM1 (leaving only the CC1α1 domain intact) did not reduce the level of CRAC current activated by store depletion (Fahrner et al., 2014), raising the possibility that the 3HB and CAD together may be able to span the gap of the ER-PM junction. Then the question becomes, does the CAD have to refold into its ‘V’ shape before it can bind to Orai1, making the CC1 coiled-coil necessary for CAD-Orai1 binding rather than for extending CAD across the junctional cleft?

Based on these results and those of prior studies, we propose the following model to describe STIM1 activation by store depletion; i.e., how the release of Ca^2+^ from the luminal domain is translated into rearrangements of the cytosolic domain that release the CAD and extend it towards the PM (**Fig. 8**). We believe the key to understanding how Ca^2+^ release is coupled to STIM1 activation comes from protein fluctuations that arise from weak intramolecular interactions. In our previous study we showed that multiple brakes operate in concert to keep STIM1 in its resting configuration (Qiu and Lewis, 2025). Because the brakes are weak, they permit thermally driven conformational fluctuations, manifested by the multiple peaks and broad distributions in most of the FRET histograms. We envision the activation trajectory as a conformational selection process involving multiple cytosolic interactions that are broken and reform along a path of lowest free energy. The transitions are stochastic and driven by thermal energy, consistent with the bidirectional coupling of luminal and cytosolic domains (**Fig. 5**). Like the trapping of STIM1 and activation of Orai1 at ER-PM junctions, the conformational activation of STIM by store depletion can be considered a type of diffusion trap but on a molecular scale. After Ca^2+^ is removed, fluctuating STIM1 domains become trapped in what is now a lower composite energy state, and a series of these trapping events ultimately leads to the active conformation. The release of Ca^2+^ from the EF hand initiates STIM1 activation by freeing the SAM domain and exposing its dimerization interface. A recent study suggests that the two SAM domains rotate in opposite directions to orient their binding domains for dimerization, and this asymmetric rotation may destabilize the CC1α1-CC3 clamp (Sallinger et al., 2024; Hogan, 2024), although the presence of flexible linker segments between the EF-SAM, TM, CC1α1, 2, and 3, and CC2 domains would likely limit the amount of torsion that can be transmitted across the ER membrane. In addition to inducing twist, dimerization also brings the TM domains closer together to destabilize the CC1 clamp and expose the CC1α1-CC3 binding interface. After transiently separating, the two CC1α2/3 hairpins can pivot down to bind their respective CC1α1 helices, effectively displacing CC1α1 from CAD and forming the 3HB intermediate, while pulling the base of CAD apart to allow its rotation out from between the two CC1 bundles. As discussed above, additional interactions with accessory proteins or membrane lipids may ultimately allow the 3HB to unfold and form a coiled-coil, while the CAD reverts to its ‘V’-shape that is now oriented towards the PM and is able to bind and activate Orai1. A full test of this hypothesis will require the isolation and structural analysis of multiple intermediate states and the ordering of transitions between them, a formidable but not insurmountable challenge.

**Figure 8.**
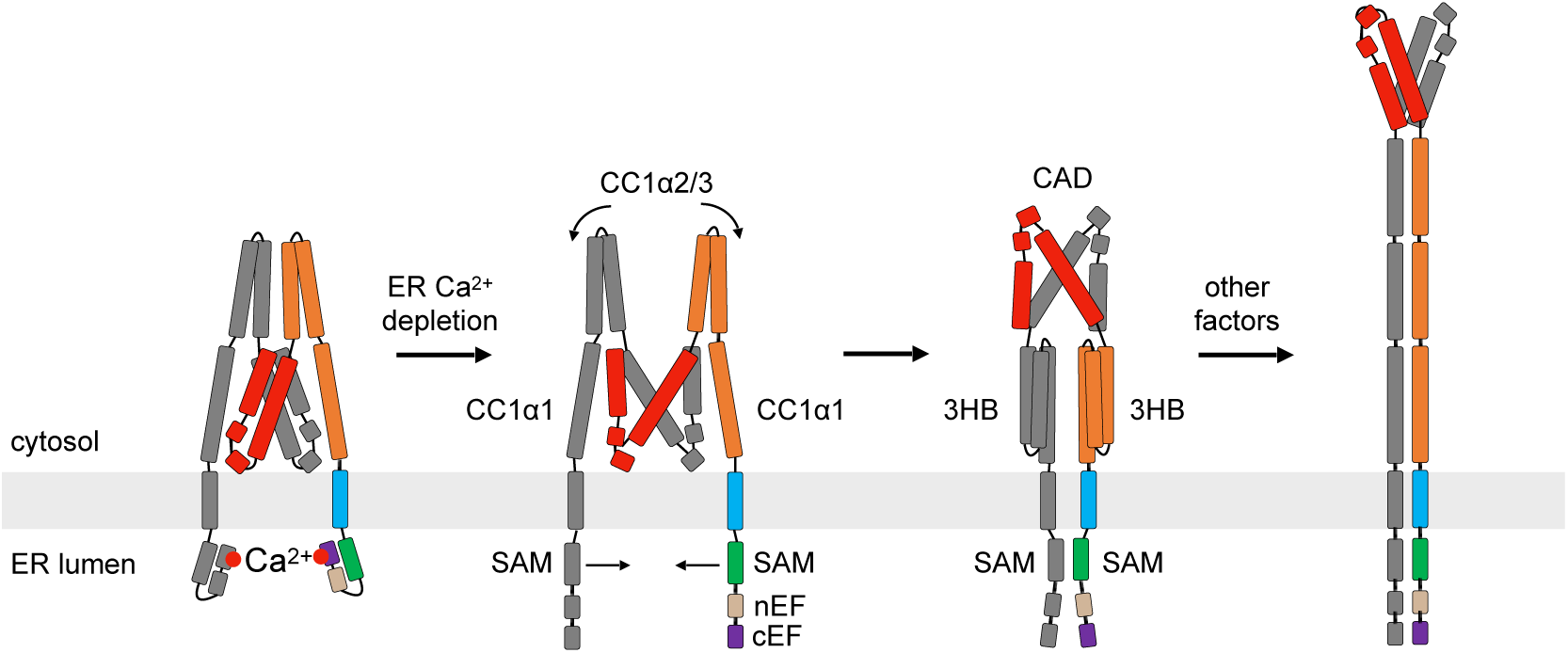
A proposed model for STIM1 activation by ER Ca^2+^ depletion. In this model, a number of events occur after ER Ca^2+^ depletion, enabled by thermally-driven structural fluctuations. The SAM domains dimerize, CC1α2/3 brake is released and rotates down to pair with CC1α1 to form a 3HB, while the CAD is rearranged to allow its escape from the CC1 clamp. Full progression to the TM-CC1 coiled-coil state occurs in the native cellular environment, enabled by cellular factors missing from the in vitro reconstituted system (see text).

## METHODS

### Cell culture

HEK 293 cells (ATCC) and HEK 293 Orai1/2/3 knockout cells (kindly provided by Dr. Mohamed Trebak, Univ. Pittsburgh) were cultured in DMEM containing 2 mM L-alanyl-glutamine, 10% FBS, and 100 U/ml penicillin/streptomycin at 37°C in 5% CO_2_. Sf9 cells were cultured in ESF 921 insect cell culture medium (Expression Systems) at 27°C with constant shaking at 120 RPM.

### DNA constructs

For insect cell expression, STIM1 constructs were cloned in the pFastBac1 vector by PCR from full-length human STIM1 (Origene) as follows. To construct his_6_-STIM1, STIM1 residues 1-34 followed by a C-terminal his_6_-tag and 3C protease cleavage sequence (LEVLFQGP) were inserted between EcoRI and XhoI restriction sites, and STIM1 residues 35-685 were inserted between XhoI and KpnI restriction sites. To construct MBP-STIM1, STIM1 residues 1-34 were inserted between BamHI and EcoRI restriction sites, the MBP sequence with a C-terminal 3C protease cleavage sequence was inserted between EcoRI and XhoI restriction sites, and STIM1 residues 35-685 were inserted between XhoI and KpnI restriction sites. All native cysteines in STIM1 (C49, C56, C227 and C437) were mutated to serines, and desired mutations were introduced by site-directed mutagenesis (Quickchange XL; Stratagene).

For puncta imaging and cysteine crosslinking experiments in HEK 293 cells, the cysteineless STIM1 and other STIM1 mutants were generated from mCherry-STIM1 (Wu et al., 2006) by site-directed mutagenesis (Quickchange XL; Stratagene). C437, the only cysteine in the cytosolic domain, was mutated to serine and the desired residues were mutated to cysteine for diamide-induced crosslinking. All plasmids were verified by Sanger sequencing or whole plasmid sequencing.

### Protein expression and purification

All STIM1 constructs in pFastBac1 vector were expressed in Sf9 cells infected with recombinant baculovirus (Bac-to-Bac, Invitrogen). For symmetric inter-subunit smFRET samples, baculovirus of His_6_-tagged STIM1 with a single cysteine substitution was used for expression. For asymmetric inter-subunit and intra-subunit smFRET samples, baculoviruses of His_6_-tagged STIM1 and MBP-tagged STIM1 were used together for heterodimer expression.

Sf9 cells were harvested 45 h post infection and were lysed in a buffer of 20 mM Tris pH 7.8, 0.5 mM EDTA, 500 µM TCEP with added protease inhibitor cocktail (Sigma-Aldrich, S8830). Cell membranes were centrifuged at 150,000 × g for 30 min at 4°C. STIM1 protein was extracted from the membranes using a buffer of 20 mM Tris pH 7.2, 150 mM NaCl, 2% n-dodecyl-β-D-maltoside (DDM), 10 mM imidazole, 500 µM tris(2-carboxyethyl)phosphine (TCEP) and benzonase (1 µl/100 ml; Sigma-Aldrich) for 1 h at 4°C. After centrifugation, Ni-NTA resin (Qiagen) was added to the supernatant and rotated for 2 h at 4°C. Ni-NTA resin was washed with 20x column volume of wash buffer containing 20 mM Tris pH 7.2, 150 mM NaCl, 0.1% DDM, 500 µM TCEP, and 40 mM imidazole. Protein was eluted in wash buffer supplemented with 300 mM imidazole. To isolate heterodimers, amylose resin (New England Biolabs) was added to the elution and rotated for 1 h at 4°C, and protein was eluted in 20 mM Tris pH 7.2, 150 mM NaCl, 0.1% DDM, 500 µM TCEP, and 10 mM maltose. To cleave the tag, 3C protease was added to the eluted protein for 1 h at room temperature. Complete cleavage was verified by SDS-PAGE. After concentrating with a 100 kDa cutoff concentrator (Amicon Ultra), the STIM1 protein was further purified by size exclusion chromatography (SEC) using a Superose 6 Increase 10/300 GL column (Cytiva) equilibrated with SEC buffer of 20 mM Tris pH 7.2, 150 mM NaCl, 0.1% DDM, and 100 µM TCEP. Fractions containing STIM1 protein were collected and concentrated with a 100 kDa cutoff concentrator (Amicon Ultra). The STIM1 protein was supplemented with 10% (v/v) glycerol and flash frozen in liquid nitrogen. The purity of STIM1 samples were higher than 90% as determined by SDS-PAGE.

His-tag MSP2N2 was obtained from Addgene (plasmid# 29520), and the S27C mutant of MSP2N2 was generated by QuickChange mutagenesis (Invitrogen) for biotin labeling. Both proteins were expressed in E. coli BL21 competent cells and purified as described (Ritchie et al., 2009).

### Protein labeling and reconstitution in liposomes

Sites for cysteine substitution and dye labeling were selected from outward-facing residues if a structure was available (EF-SAM, CAD), and if not, from residues predicted to lie outside interface surfaces. STIM1 dimer containing two cysteines was labeled with maleimide-conjugated Alexa Fluor 555 and Alexa Fluor 647 (ThermoFisher). STIM1 was diluted to 10 µM in 200 µl desalting buffer (20 mM Tris pH 7.2, 150 mM NaCl, 0.1% DDM, and 100 µM TCEP). 5 µM donor fluorophore and 5 µM acceptor fluorophore were added and after incubation for 1 h at 4°C, free dye was removed with a home-packed desalting column filled with 5 ml of G50 resin (Sigma) pre-equilibrated with desalting buffer. Labeling efficiency was ∼50% for most samples. The labeled STIM1 protein was concentrated to ∼10 µM, aliquoted, supplemented with 10% (v/v) glycerol and flash frozen.

To reconstitute STIM1 proteins in liposomes, egg PC, POPE, and POPG (Avanti) were mixed at a 3:1:1 ratio in chloroform, and 0.5% biotinylated PE (Avanti) was added. The lipids in chloroform were dried by nitrogen gas and kept in vacuum overnight. The dry lipids were resuspended in buffer containing 20 mM Tris pH 7.2 and 150 mM NaCl with 2 mM CaCl_2_ or 0.5 mM EGTA to a final concentration of 10 mg/ml, creating a cloudy solution. Using an extruder set (Avanti Polar Lipids) the lipids were then extruded through a membrane with 100-nm pores (Whatman) 30 times to form liposomes with a diameter of ∼100 nm. For each sample, 180 µl of liposomes (10 mg/ml) was added to 20 µl of 12% n-octyl-β-D-glucopyranoside (βOG) solubilized in the same buffer and rotated for 15 min at 4°C. 10 µl of labeled STIM1 protein (∼10 µM) was added to the liposomes and rotated for 30 min at 4°C. 25 mg of Bio-Beads (Bio-Rad) was added to the sample and rotated for 1 h at 4°C, and another 25 mg of Bio-Beads was added and rotated for 1 additional hour. The reconstituted proteoliposomes were separated from free detergent and free STIM1 protein using a home-packed Sepharose CL-4B column (Sigma Aldrich). The fraction containing STIM1 proteoliposomes was added to 10 mg Bio-Beads and rotated overnight at 4°C to remove residual detergent, at which point the sample was ready for smFRET experiments.

For experiments using nanodiscs (**Figs. 4C-E and 7**), S27C MSP2N2 was labeled with maleimide-PEG11-biotin (Thermo Fisher #21911) as follows. S27C MSP2N2 was diluted to 50 µM in 200 µl TBS buffer (20 mM Tris pH 7.2, 150 mM NaCl, 100 µM TCEP). 200 µM maleimide-PEG11-biotin was added and after incubation for 2 h at 4°C, free maleimide-PEG11-biotin was removed with a home-packed desalting column filled with 5 ml of G50 resin (Sigma) pre-equilibrated with TBS buffer. To reconstitute STIM1 proteins into MSP2N2 nanodiscs, dye-labeled STIM1 protein, MSP2N2 and lipids were mixed at a molar ratio of 1:2:200 in buffer containing 20 mM Tris pH 7.2, 150 mM NaCl, and 20 mM Na cholate. The final concentration of STIM1 monomer in 1 ml of reaction volume was 1 µM. The lipid mixture contained PC, PE, and PG at a ratio 3:1:1. To avoid possible constraint of STIM1 movement caused by double biotin-streptavidin binding of a single nanodisc, biotin-labled S27C and unlabled WT MSP2N2 were mixed at a ratio of 1:5 to minimize the double-biotin nanodiscs while producing enough single-biotin nanodiscs.

The mixture was rotated for 30 min at 4°C. 250 mg of Bio-Beads (Bio-Rad) was added to the sample and rotated for 1 h at 4°C, and another 250 mg of Bio-Beads was added and rotated overnight. The reconstituted nanodiscs were separated from empty nanodiscs by size exclusion chromatography (SEC) using a Superose 6 Increase 10/300 GL column (Cytiva) equilibrated with SEC buffer containing 20 mM Tris pH 7.2, 150 mM NaCl, and the eluted sample was ready for smFRET experiments.

### Imaging chamber preparation

Imaging chambers for single-molecule fluorescence imaging were prepared according to established protocols (Roy et al., 2008). Briefly, microscope slides and coverslips were first cleaned by stepwise sonication for 15 min each in glass containers with acetone, ethanol, 1 M KOH and pure water. They were then coated with a 100:1 PEG/PEG-biotin mixture (Laysan Bio) prior to flow cell construction. Strips of double-sided tape were applied to a quartz microscope slide (Finkenbeiner) to form channel walls, with holes at both ends of each channel to allow exchange of sample solutions. A microscope coverslip of 1.5 thickness (Erie Scientific) was pressed on the tape strips and edges of the channels were sealed with epoxy glue (Devcon).

Prior to attaching biotinylated liposomes to the coverslip surface, channels were rinsed with 20/150 TBS (150 mM NaCl, 20 mM Tris, pH 7.2 with HCl) then incubated with 0.2 mg/ml neutravidin (Thermo Fisher CWA) for 5 min. Liposomes were loaded into the channels after washing out the neutravidin. Prior to TIRF imaging, channels were filled with imaging buffer (20 mM Tris pH 7.2, 150 mM NaCl, 1 % D-glucose, 100 μM cyclooctatetraene, 1 mg/ml glucose oxidase (Sigma Aldrich), and 0.04 mg/ml catalase) containing either 2 mM CaCl_2_ or 0.5 mM EGTA.

### TIRF microscopy and smFRET measurements

All smFRET experiments were performed at room temperature following a previous protocol (van Dorp et al., 2021) with some modifications. The home-built TIRF system is based on an Axiovert S100 TV microscope equipped with a Fluar 100x 1.45 NA oil-immersion objective (Zeiss). 532– and 637-nm lasers (OBIS 532 nm LS 150 mW, Coherent and OBIS 637 nm LX 140 mW, Coherent) were used for excitation in objective TIRF mode. Donor and acceptor signals were separated by a 652 nm dichroic (Semrock) and passed through 580/60 nm and 731/137 nm bandpass filters (Semrock), respectively, mounted in an OptoSplit-II beamsplitter (Cairn Research) to an EM-CCD camera (iXon DU897E, Andor). Hardware and data acquisition were controlled by homemade scripts in μManager as described (van Dorp et al., 2021).

For each sample, the molecule density and distribution on the coverslip surface was checked with the TIRF microscope. When optimal density was achieved (300 to 500 molecules in the camera field of view), donor and acceptor emission were recorded with a 100-ms integration time under 532-nm laser excitation for 80 s, immediately followed by 1 s of 637-nm laser excitation to verify acceptor dye bleaching.

Data were analyzed using custom Python scripts as detailed previously (van Dorp et al., 2021). Briefly, molecules with single acceptor bleaching before single donor bleaching were selected. The FRET ratio E was calculated at each time point as E = I_A_/(I_A_ + γI_D_), where I_A_ and I_D_ are the acceptor and donor fluorescence values, respectively, and γ was measured empirically for each molecule as described previously (van Dorp et al., 2021). For single-molecule traces longer than 20 points (2 s), summed FRET histograms were constructed by distributing FRET amplitudes into 40 bins (from –0.25 to 1.25) and normalizing by the number of points in each trace. Histograms were fitted with a sum of Gaussian functions using routines written in Igor Pro (WaveMetrics).

R_0_ of the dye pair was measured empirically on our smFRET system. The crystal structure of CAD was used to simulate the dye-dye distances for 417-417’ and 431-431’ pairs using Crystallography and NMR System (CNS) (Brunger, 2007). These distances were used to solve for R_0_ using the FRET equation and 417:417’ and 431:431’ smFRET measurements from ctSTIM1 samples. The calculated R_0_ values were 5.85 nm for 417:417’ and 5.80 nm for 431:431’; a value of 5.8 nm was used to estimate distances throughout this paper. Distances (R) were calculated from FRET (E) using the relation E = 1/[1 + (R/R_0_)^6^].

### Cysteine-crosslinking smFRET experiments

To test the effect of S339C and Y361C crosslinking on STIM1 activation, smFRET measurements were performed with STIM1 nanodisc samples which allowed full exposure of STIM1 to buffer exchange in the chamber. The molecule density on the coverslip surface was optimized to a level of 300-500 molecules in the TIRF microscope field of view. To crosslink the cysteine pair (S339C or Y361C), a buffer of 20 mM Tris pH 7.2, 150 mM NaCl, 2 mM CaCl_2_ and 1 mM diamide was injected into the chamber. After 10 min, the solution was exchanged with imaging buffer containing 2 mM Ca^2+^ or 0.5 mM EGTA for smFRET measurements. After measurements of the crosslinked condition, EGTA buffer with 50 mM DTT was injected into the chamber to break the disulfides, and after 10 min the this solution was exchanged with imaging buffer containing 0.5 mM EGTA and 10 mM DTT for smFRET measurement.

### Dye position simulation with Crystallography and NMR System (CNS)

We simulated dye positions on STIM1 structures using Crystallography and NMR System (CNS) as described (Brunger, 2007). Briefly, the pdb file was loaded in Pymol and residues for dye labeling were mutated to cysteine. Atomic models of the fluorophores and their maleimide linker were attached to the mutated cysteines and subjected to molecular dynamics simulations by fixing all protein atoms except the dye and linker atoms. 100 simulations were performed for each labeling pair and the average distance between the dye centers (CAO atom) was calculated from the resulting coordinates.

### STIM1 puncta imaging

Orai1/2/3 knock-out HEK 293 cells were transfected with mCherry-flSTIM1 constructs with WT, S339C or Y361C mutants. 24 h after transfection, cells were trypsinized and plated on matrigel treated glass coverslips overnight. Imaging was conducted about 48 h after transfection on a Nikon Eclipse Ti2 microscope equipped with an Apo TIRF 100x 1.49 NA oil objective. Cells were excited with 561 nm laser under TIRF mode, and emission that passed through 600 ± 50 nm band pass filter was collected by a Photometrics prime 95B camera. Before the imaging acquisition started, the cells were changed into standard 2 Ca Ringer’s solution contained (in mM): 155 NaCl, 4.5 KCl, 2 CaCl_2_, 1 MgCl_2_, 10 D-glucose and 5 Na-HEPES (pH 7.4). 2 Ca Ringer’s solution with 1 mM diamide or with 1 mM diamide + 1 μM thapsigargin was applied to the cells at times indicated in Figs. 4 and 7.

Images were analyzed with ImageJ. After background correction, the mean intensity of the cell footprint was measured for each frame. The time-resolved puncta intensity was normalized by the first frame for each cell before calculating the mean and SEM, in order to compensate for differences in STIM1 expression.

### Cysteine crosslinking and Western blot analysis

For crosslinking flSTIM1 in situ, HEK 293 cells were transfected with mCh-STIM1 constructs with S339C or Y361C mutants. 48 h after transfection, cells were exposed to 0.5 mM diamide for 10 min in Ringer’s solution containing 2 mM Ca^2+^. Cells were then lysed in RIPA buffer containing 20 mM NEM, 0.5 mM EDTA, and protease inhibitor cocktail (Cell Signaling Technology). Samples were run on SDS-PAGE and analyzed by Western blot using rabbit anti-mCherry antibody (1:2000, OriGene Technologies), with secondary antibodies 680RD anti-rabbit on a LI-COR Odyssey imaging system. Western blots were scanned and analyzed using ImageJ, and the area under each peak indicated the amount of protein in each band. Crosslinking efficiencies were calculated by dividing the amount of STIM1 in dimer band (crosslinked) by the total amount of STIM1 (monomer + dimer).

## ACKNOWLEDGEMENTS

The authors thank Dr. Mohamed Trebak (Univ. of Pittsburgh) for providing the Orai1/2/3 triple-knockout HEK 293 cell line, and members of the Lewis lab for helpful discussions during the course of this study. This work was supported by NIH grant R37GM45374, R35GM149305, and the Mathers Charitable Foundation (RSL).

## Author Contributions

RQ: designed research, performed research, analyzed data, wrote the paper. RSL: designed research, analyzed data, wrote the paper.

## Classification

Biological Sciences / Physiology

## SUPPLEMENTARY FIGURES

**Figure S1.**
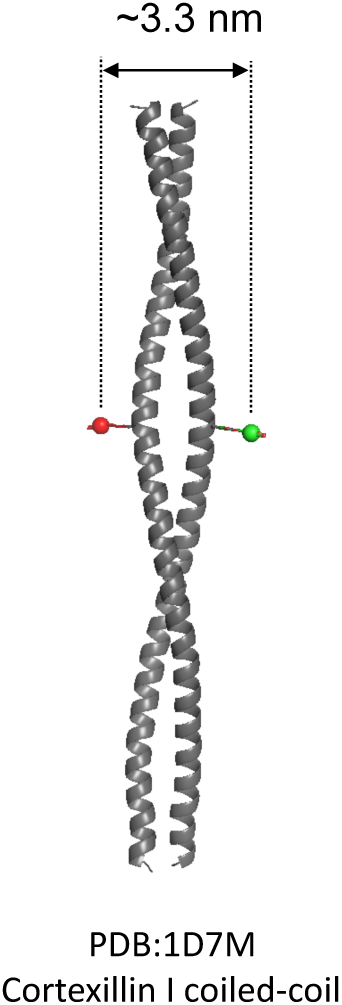
Dye-dye distance simulation in cortexillin I coiled-coil structure. Dye positions were simulated using a CNS-based approach based on the cortexillin I coiled-coil structure (PDB: 1D7M). The inter-dye distance across the two symmetric helices of the coiled-coil was estimated to be approximately 3.3 nm.

**Figure S2.**
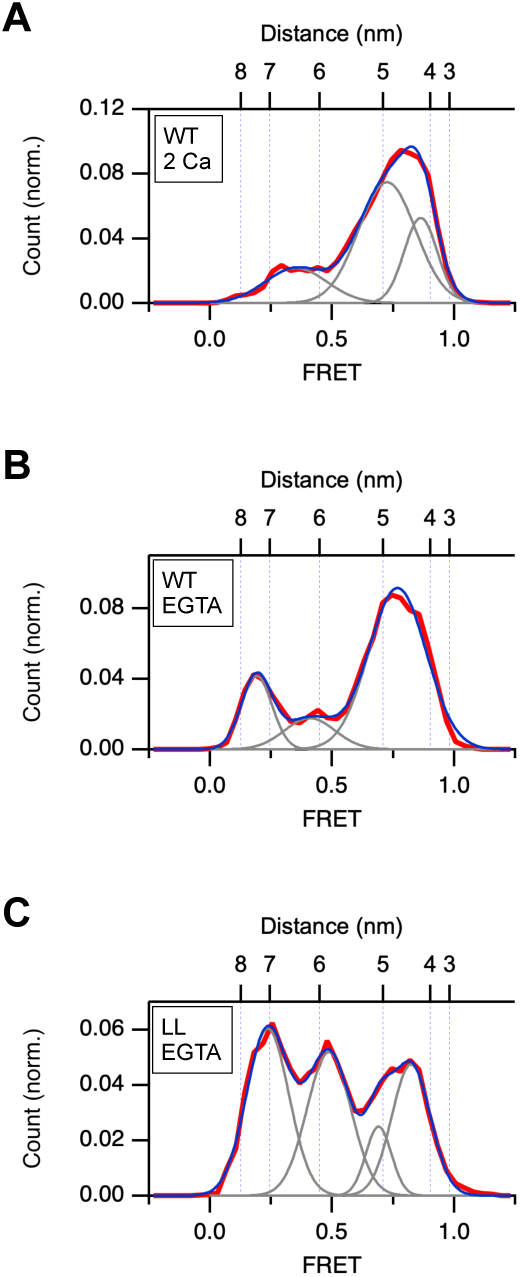
Effects of the N234L/S237L double mutation (LL) on the CC1α2/3 hairpin. CC1α2-CC1α3 (291:325) FRET histograms are shown with fitted Gaussian curves (gray) and their sum (blue) superimposed on the data (red). (*A*) WT STIM1 in 2 mM Ca^2+^ (n=202). Fit parameters (peak FRET and fractional area): 0.35 (18%), 0.73 (58%), 0.86 (24%). (*B*) WT STIM1 in 0.5 mM EGTA (n=246). Fit parameters (peak FRET and fractional area): 0.19 (18%), 0.42 (12%), 0.75 (60%), 0.88 (10%). (*C*) STIM1-LL in 0.5 mM EGTA (n=219). Fit parameters (peak FRET and fractional area): 0.24 (35%), 0.49 (31%), 0.69 (9%), 0.82 (25%). The significant increase of low FRET fraction in STIM1-LL indicates the unfolding and extension of the CC1α2/3 hairpin, consistent with the formation of the CC1 coiled-coil.

**Figure S3.**
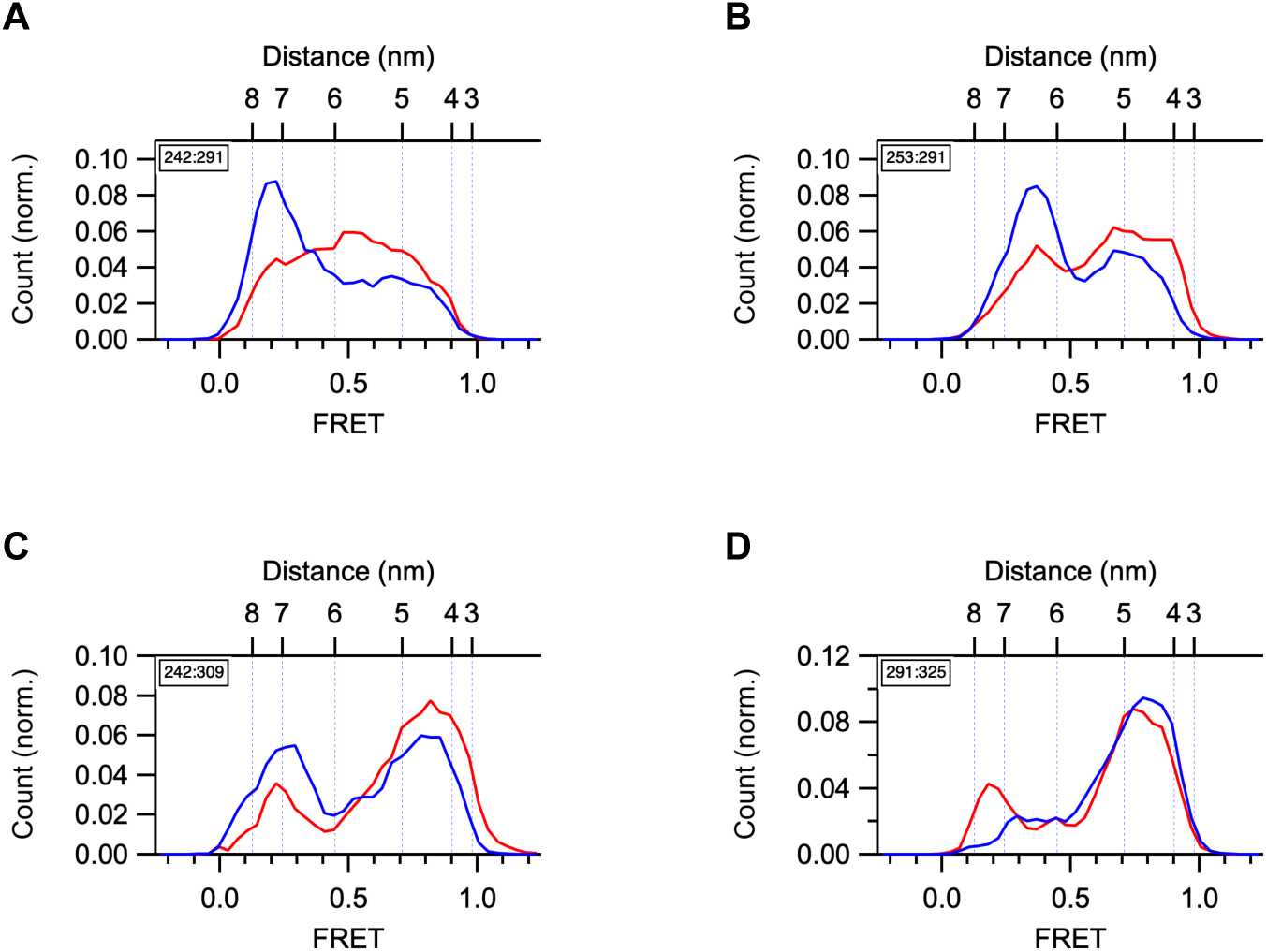
Comparison of intrasubunit distances in CC1 in Ca^2+^ and EGTA. FRET histograms are shown for 2 mM Ca^2+^ (blue) and 0.5 mM EGTA (red) with numbers of molecules (in Ca^2+^, in EGTA). (*A*) 242:291 (n=192, 189). (*B*) 253:291 (n=254, 275). (*C*) 242:309 (n=82, 158). (*D*) 291:325 (n=202, 210).

**Figure S4.**
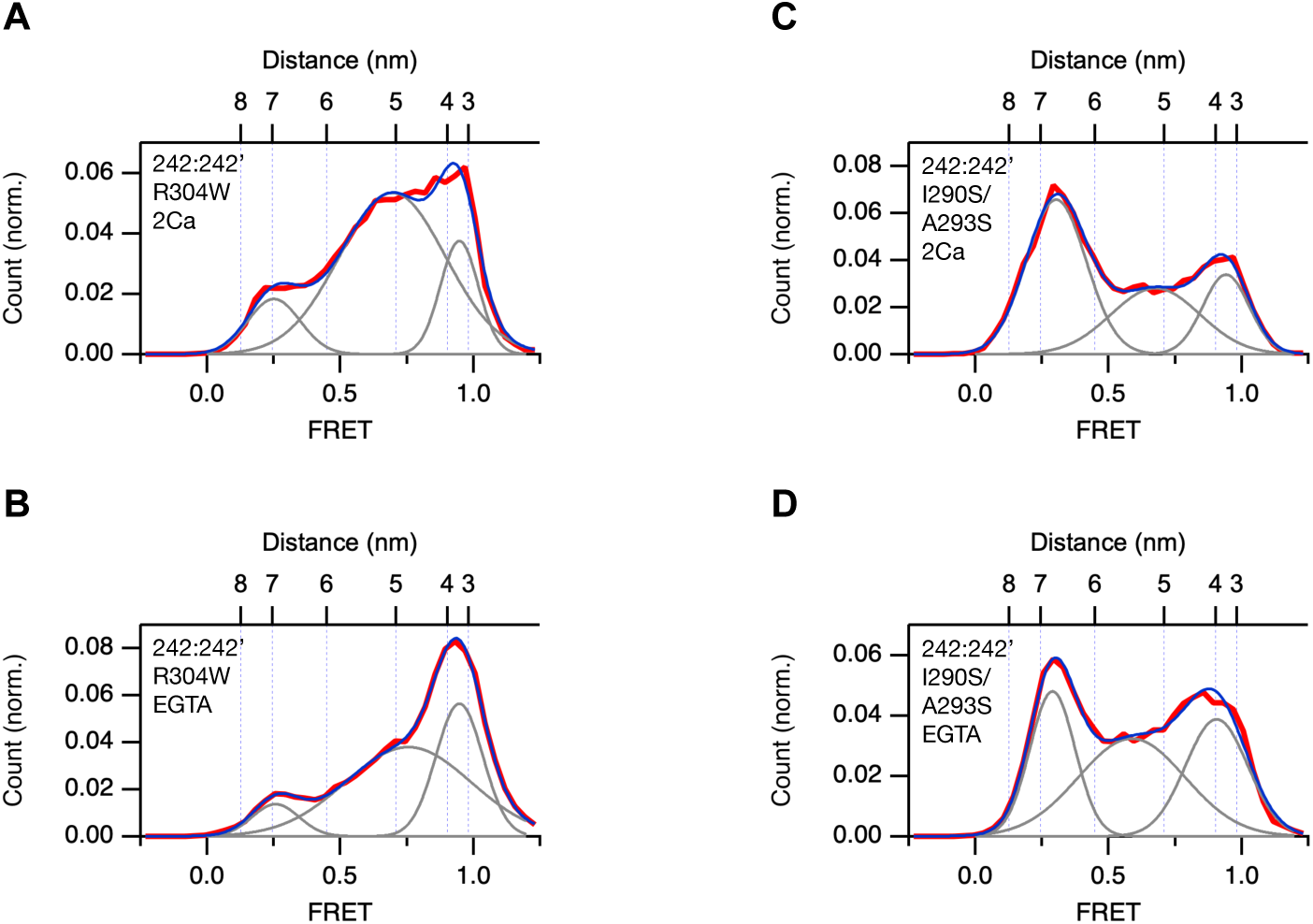
Destabilizing the 3HB promotes transitions to the fully extended coiled-coil state. smFRET histograms are shown with fitted Gaussian curves (gray) and their sum (blue) superimposed on the data (red). (*A*) CC1α1-CC1α1’ FRET histogram of STIM1-R304W in 2 mM Ca^2+^ (242:242’; n=225). Fit parameters (peak FRET and fractional area): 0.25 (12%), 0.70 (70%), 0.95 (18%). (*B*) CC1α1-CC1α1’ FRET histogram of STIM1-R304W in 0.5 mM EGTA (242:242’; n=356). Fit parameters (peak FRET and fractional area): 0.26 (8%), 0.76 (59%), 0.95 (33%). (*C*) CC1α1-CC1α1’ FRET histogram of I290S/A293S-STIM1 in 2 mM Ca^2+^ (242:242’; n=316). Fit parameters (peak FRET and fractional area): 0.30 (49%), 0.68 (31%), 0.94 (20%). (*D*) CC1α1-CC1α1’ FRET histogram of STIM1-I290S/A293S in 0.5 mM EGTA (242:242’; n=434). Fit parameters (peak FRET and fractional area): 0.29 (27%), 0.59 (43%), 0.91 (30%).

**Figure S5.**
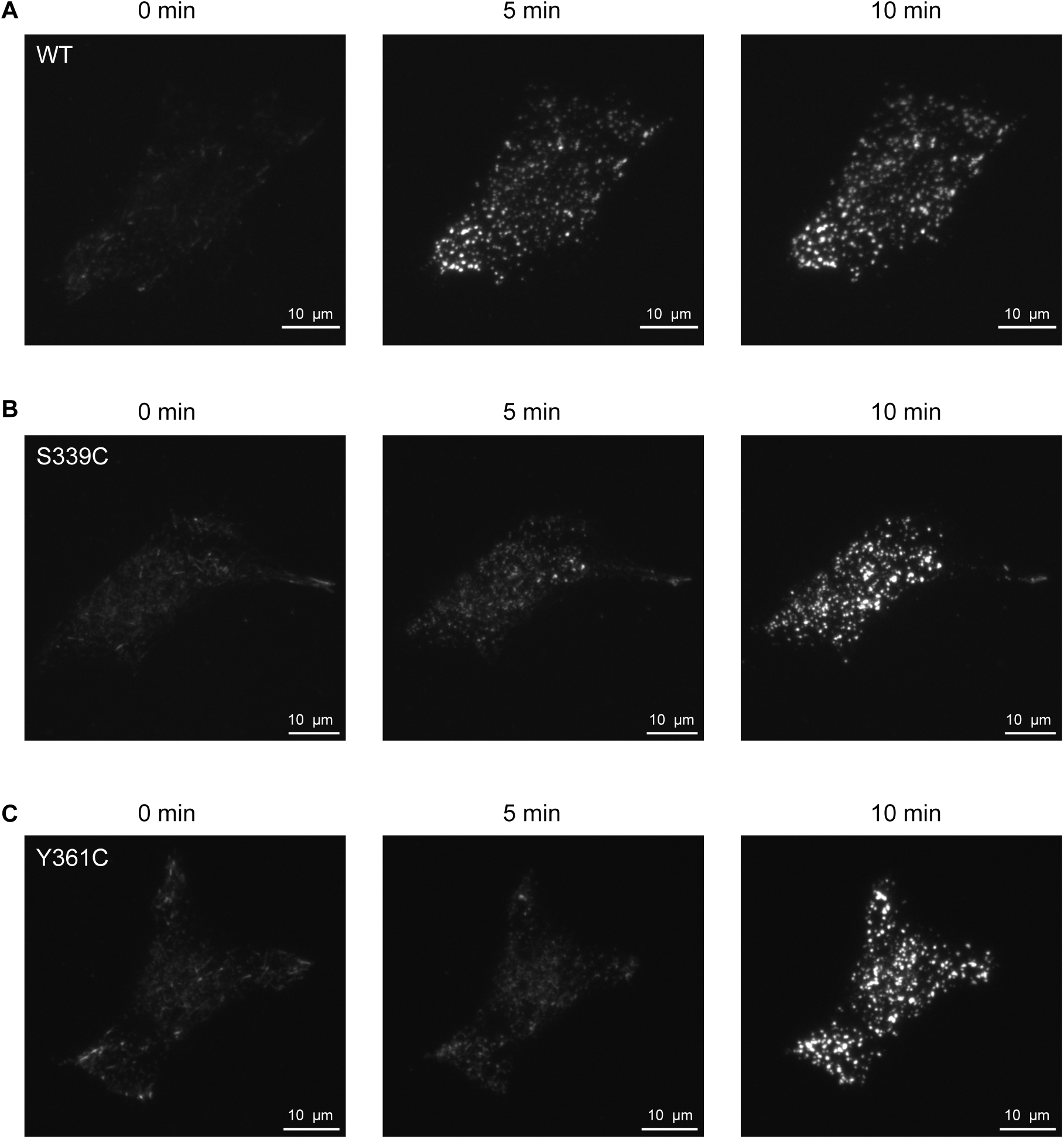
Kinetics of STIM1 puncta formation are slowed by crosslinking S339C or Y361C. Representative TIRF images of mCh-STIM1 puncta at different time points for STIM1-WT (*A*), STIM1-S339C (*B*), and STIM1-Y361C (*C*), taken from the experiments shown in **Figs. 4 and 7**.

## SUPPLEMENTARY TABLES

**Table S1.** Comparison of FRET-derived distances in 2 mM Ca^2+^ with the CAD crystal structure and the AlphaFold2 model of STIM1. For each dye pair, the predominant smFRET value from the STIM1-WT amplitude histograms is listed and used to calculate a distance. The number of single molecule traces used to construct each histogram is indicated. Inter-dye distances from the CAD crystal structure (3TEQ.pdb) and AlphaFold2 model (Qiu and Lewis, 2025) were measured as described in Methods. NA indicates that the residues are absent in the CAD crystal structure.

| Dye pair | Peak FRET | Number of molecules | Distance from FRET (nm) | Distance from CAD crystal structure (nm) | Distance from AlphaFold2 (nm) |
| --- | --- | --- | --- | --- | --- |
| 337:337' | 0.78 | 183 | 4.70 | NA | 4.08 |
| 349:349' | 0.84 | 221 | 4.40 | 4.29 | 4.31 |
| 378:378' | 0.93 | 226 | 3.77 | 3.51 | 3.41 |
| 431:431' | 0.78 | 204 | 4.70 | 4.69 | 4.85 |
| 349:431 | 0.86 | 223 | 4.29 | 3.88 | 4.12 |
| 349:431' | 0.97 | 232 | 3.25 | 3.17 | 2.74 |

